# A sensitive assay for dNTPs based on long synthetic oligonucleotides, EvaGreen dye, and inhibitor-resistant high-fidelity DNA polymerase

**DOI:** 10.1101/2019.12.17.879122

**Authors:** Janne Purhonen, Rishi Banerjee, Allison E McDonald, Vineta Fellman, Jukka Kallijärvi

## Abstract

Deoxyribonucleotide triphosphates (dNTPs) are vital for the biosynthesis and repair of DNA. Their cellular concentration peaks during the S phase of the cell cycle. In non-proliferating cells dNTP concentrations are low, making their reliable quantification from tissue samples of heterogeneous cellular composition challenging. Partly because of this, the current knowledge related to regulation of and disturbances in cellular dNTP concentrations derive from cell culture experiments with little corroboration at the tissue or organismal level. Here, we fill the methodological gap by presenting a simple non-radioactive microplate assay for the quantification of dNTPs with a minimum requirement of 10 to 30 mg of biopsy material. In contrast to published assays, this assay is based on long (~200 nucleotides) synthetic single-stranded DNA templates, an inhibitor-resistant high-fidelity DNA polymerase, and the double-stranded-DNA-binding EvaGreen dye. The assay quantifies reliably as little as 100 fmol of each of the four dNTPs. Importantly, the assay allowed measurement of minute dNTP concentrations in mouse liver, heart, and skeletal muscle.

## INTRODUCTION

Deoxyribonucleotide triphosphates (dNTPs) are building blocks of DNA, and accordingly their cellular concentration is highest during active DNA synthesis, such as in proliferating cells (1). Basal dNTP levels in non-proliferating or postmitotic cells are only approximately 5% that of in cells in S phase and are mainly required for DNA repair and maintenance of mitochondrial DNA (1, 2). While studying pyrimidine biosynthesis in a mouse model of mitochondrial respiratory complex III deficiency (3), we encountered the problem that no suitable method, applicable for the general laboratory, to measure dNTPs from tissue samples exists. Traditional enzymatic methods to measure dNTPs have relied on the incorporation of radioactive dNTPs to a complementary strand of a DNA template by a DNA polymerase (4, 5). These methods have mainly allowed measurement of dNTPs from actively proliferating cultured cells with high dNTP concentrations and limited complexity of interfering biological matrix. Recently, a HPLC-MS-based method was developed that allows quantification of tissue dNTP levels from a feasibly small tissue amount (15 to 30 mg) (6). However, not every laboratory has suitable HPLC-MS equipment and expertise. A simple probe hydrolysis-based enzymatic fluorescence assay has also been developed to measure dNTPs from cultured cells (7). This assay utilizes synthetic DNA templates with a 3’ primer-binding region, mid-template dNTP-detection region, 5’ probe-binding sequence, and a Taq DNA polymerase with 3’ to 5’ exonuclease activity to hydrolyse the probe - an adaptation of typical TaqMan qPCR chemistry. The authors achieved low baseline fluorescence by using dual-quenched 6-carboxyfluorescein (FAM) - labelled probes. Due to the simplicity of the assay, we set out to test this assay as a means to quantify dNTPs in mouse liver extracts. We found some limitations in the original assay, especially concerning complex tissue extracts with low dNTP concentrations, and developed a novel even simpler assay based on a long synthetic template, EvaGreen DNA dye, and a robust inhibitor-resistant high-fidelity DNA polymerase.

## MATERIALS AND METHODS

### Reagents

Integrated DNA Technologies (Coralville, Iowa, USA) synthetized all DNA oligonucleotides (Supplementary Table 1). We used standard desalted DNA oligonucleotide preparations for the novel assay and PAGE-purified primers and templates for the published (7) probe-hydrolysis based assay. The probes were HPLC purified. All critical reagents, materials, and instrumentation are listed in Supplementary Table 2.

### Molecular cloning

Alternative oxidase (AOX) cDNA from Pacific oyster (*Crassostrea gigas*) (Genbank, #ACL31211) (8) was cloned into the pcDNA6A mammalian expression vector (Invitrogen). First, an internal EcoRI site was removed by PCR mutagenesis (primers 5’-ATT CGC ACC AGC AAT GGG CT-3’ and 5’-ACC ACT GAC CCT CAG TAG AAT T-3’) and then the coding sequence was PCR-amplified (primers 5’-ATGAATTC ATG GGA AGT TTG CGA CAA ATA AC-3’and 5’-ATCTCGAG TCA CTT CCC TGG CTC ATA AGG-3’ or 5’-ATCTCGAG CTT CCC TGG CTC ATA AGG ATT-3’) and cloned into pcDNA6A with or without a C-terminal V5-6XHis tag. The cDNA was further subcloned into the Sleeping Beauty transposon (9) donor plasmid ITR\CAG-MCS-IRES-Puro2A-Thy1.1\ITR (a kind gift from Dr. Madis Jakobson, Max Planck Institute of Biochemistry, Germany).

### Cell culture, transfection and immunofluorescence staining

Mammalian cells were cultured in standard high-glucose medium (DMEM, 10% serum, Glutamax, penicillin and streptomycin). The expression and subcellular localization of the Pacific Oyster AOX (PoAOX) was confirmed by transfecting the pcDNA6-poAOX-V5-6XHis construct into COS-1 fibroblast-like cells derived from *Cercopithecus aethiops* kidney, followed by immunofluorescence staining (Supplementary Figure 1A). To visualize mitochondria and PoAOX, mouse anti-MT-CO1 (ab14705, Abcam) and rabbit anti-V5 (AB3792, Merck) antibodies were used, respectively.

To generate a cell line stably expressing PoAOX, mouse hepatoma cells (Hepa1-6) were co-transfected with a plasmid encoding SB100X transposase and the Sleeping Beauty transposon donor plasmid carrying the PoAOX-V5-6XHis insert (1:10 ratio) using FuGENE® HD transfection reagent. For a control cell line, the empty ITR\CAG-MCS-IRES-Puro2A-Thy1.1\ITR plasmid was transfected. After puromycin selection (2 µg/ml), resistant colonies were pooled, and after 5 passages the cell were cultured without puromycin. The activation and activity of poAOX was assessed by measuring antimycin A-resistant respiration using Oxygraph-2k (OROBOROS instruments) (Supplementary Figure 1B).

For dNTPs measurements, the cells were plated into T75 flasks (ThermoFisher Scientific). The next day, either vehicle or 5 µM 5-fluorouracil (from 5 mM stock in H_2_O) or 200 nM myxothiazol (from 0.5 mM stock in EtOH) was added. After approximately 6 hours the cells were washed with phosphate-buffered saline (PBS) and detached with TrypLE (ThermoFisher Scientific). Then, 10 ml ice-cold PBS was added, a 0.1-ml aliquot was saved for counting of the cells, and the cells were pelleted by centrifugation (300g for 5 min.) at +4°C. The cell pellets were resuspended in 550 µl of ice-cold 60% MeOH and stored at −80°C until extraction. To obtain samples with a decreased number of proliferating cells, the cells were allowed to grow to near 100% confluence and then processed as described above.

### Collection and homogenization of mouse tissues

Mice of congenic C57BL/6JCtrl background were maintained in individually-ventilated cages at the animal facilities of University of Helsinki, Finland, under the internal license KEK19-033. Littermate 110-day old male mice were euthanized by cervical dislocation and the tissues were immediately excised and placed in liquid nitrogen and stored at −80°C. The frozen tissue samples were directly homogenized in ice-cold 60% MeOH (typically, 40 mg/0.55 ml) using battery-operated microtube pestles for liver and roughened glass-to-glass tissue grinders for heart and skeletal muscle (whole calf).

### Extraction of dNTPs

The cultured cells and mouse tissue homogenates in 60% MeOH were further denatured by incubation at 95°C for 3 min to quench residual enzymatic activity and to facilitate the extraction. Supernatants (550 µl) from an 18500g centrifugation (6 min at +4°C) were run through 3-kDa cut-off centrifugal filters (Amicon Ultra-0.5 ml, Merck) into pre-weighted collection tubes to remove remaining macromolecules. Next, MeOH and hydrophobic metabolites were removed by washing the extracts twice with 1.4 ml diethyl ether. Residual diethyl ether was evaporated using Speed-Vac Plus SC110A centrifugal vacuum evaporator (Savant Instruments, Farmingdale, NY, USA) (typically 15 min at 65°C). The remaining liquid after evaporation was estimated by weighing the tubes and assuming that 1 µl weights 1 mg. Final tissue extract volumes were adjusted to 80-160 µl per 40 mg of initial tissue weight (160 µl/40 mg was considered optimal). Extracts from proliferating cultured cells were adjusted to 100-400 µl per 10^6^ cells. If the extracts were not immediately subjected to measurements, they were stored at −80°C until use.

### Probe hydrolysis-based enzymatic assay for dNTPs

For the fluorochrome-quencher-probe -based polymerase assay for dNTPs, we followed the published protocol (7) with minor modifications. To accommodate the relatively higher sample volume and 384-well format, we lowered the reaction volume from 25 to 10 µl and added the reaction components as a 2x-master mix followed by addition of an equal volume of sample. The final reaction concentrations were: 0.4 µM primers, probes and templates, 100 µM non-limiting dNTPs, 2 mM MgCl_2_, and AmpliTaq Gold DNA polymerase at 35 U/ml. The reaction was started by activating the hot-start polymerase by a 10-minute incubation step at 95°C. Baseline fluorescence was read at 60°C immediately after polymerase activation. Thereafter, the reaction was allowed to proceed at 60°C while monitoring the fluorescence. The Bio-Rad CFX384 qPCR instrument served as a thermal cycler and fluorometer for the assay.

### Q5 DNA polymerase and EvaGreen-based assay for dNTPs

The following reaction concentrations were found to be optimal: 1x Q5 reaction buffer: 0.25 µM primer and 0.2 µM template, 25 µM non-limiting dNTPs, 1x EvaGreen (Biotum) and 20 U/ml Q5 DNA polymerase (New England Biolabs). The reaction components were prepared as a 2x-master mix and 5 µl of the mix was pipetted into opaque wells of 384-well PCR-plate (Bio-Rad). Then an equal volume of sample was added. The reaction set-up was prepared on ice. The thermal cycler (CFX384, BioRad) was programmed to heat the plate to 98°C (10s) followed by cooling to the final reaction temperature (67°C was considered optimal). The baseline fluorescence was immediately read after reaching the target temperature. Thereafter the fluorescence (SYBR Green / FAM channel of the instrument) was recorded typically once every 5 minutes for 75 minutes. To export the raw fluorescence values for data analysis, the automatic baseline correction by the instrument was turned off. A step-by-step protocol is available as a Supplementary File.

### Data analysis

Second-order polynomial curve fits using the least squares regression were calculated using GraphPad Prism 8.2.1 (GraphPad Software, CA, USA). Signal-to-noise ratio was defined as background-subtracted fluorescence (n=3) divided by the standard deviation of the background (n=6). The lowest limit of quantification (LLOQ) was estimated as the lowest concentration at which 33% differences could still be reliably distinguished. At this concentration signal-to-noise ratio was equal to or higher than 10. The lowest limit of detection was defined as the lowest concentration at which the signal-to-noise ratio was higher than 2. The measured concentrations were corrected for losses during 3 kDa cut-off filtration. This was accomplished by weighing the residual liquid in the filter device and assuming a 0.92 g/ml density of 60% MeOH on ice.

## RESULTS

### Design of the assay concept

We initially tested the published probe hydrolysis-based enzymatic assay (7) for measurement of hepatic dNTPs. However, we encountered two problems. First, the suggested AmpliTaq Gold DNA polymerase proved to be sensitive to inhibition by liver extracts (Figure 1, Supplementary Figure 2). Second, we observed a time-dependent increase in background fluorescence, which greatly limited the sensitivity (Figure 1A). We reasoned that the increasing background fluorescence could originate from the low-fidelity nature of the non-proofreading polymerase leading to eventual incorporation of a wrong (non-complementary) deoxyribonucleoside in the presence of the highly unbalanced dNTP concentrations required by the assay. Given that no commercially available true high-fidelity polymerase with 5’ to 3’ exonuclease activity required for probe hydrolysis exists, we replaced the fluorochrome- and quencher-labelled probes with EvaGreen double-stranded DNA (dsDNA) dye, increased template length to 197 nucleotides, and performed the assay in the presence of an inhibitor-resistant high-fidelity Q5 DNA polymerase (Figure 1B, D and Figure. 2). The designed template DNA oligomers had the same primer-binding and a single dNTP-detection site at the 3’end as in the original probe-based assay (Supplementary Table 1). The EvaGreen-based detection system produced an increasing fluorescence signal, which was proportional to the added limiting dNTP (Figure 1B). Most importantly, the background fluorescence remained nearly constant during the whole reaction time, and the liver extracts spiked with known amounts of dNTPs also produced the expected increase in signal (Figure 1D).

**Figure 1.**
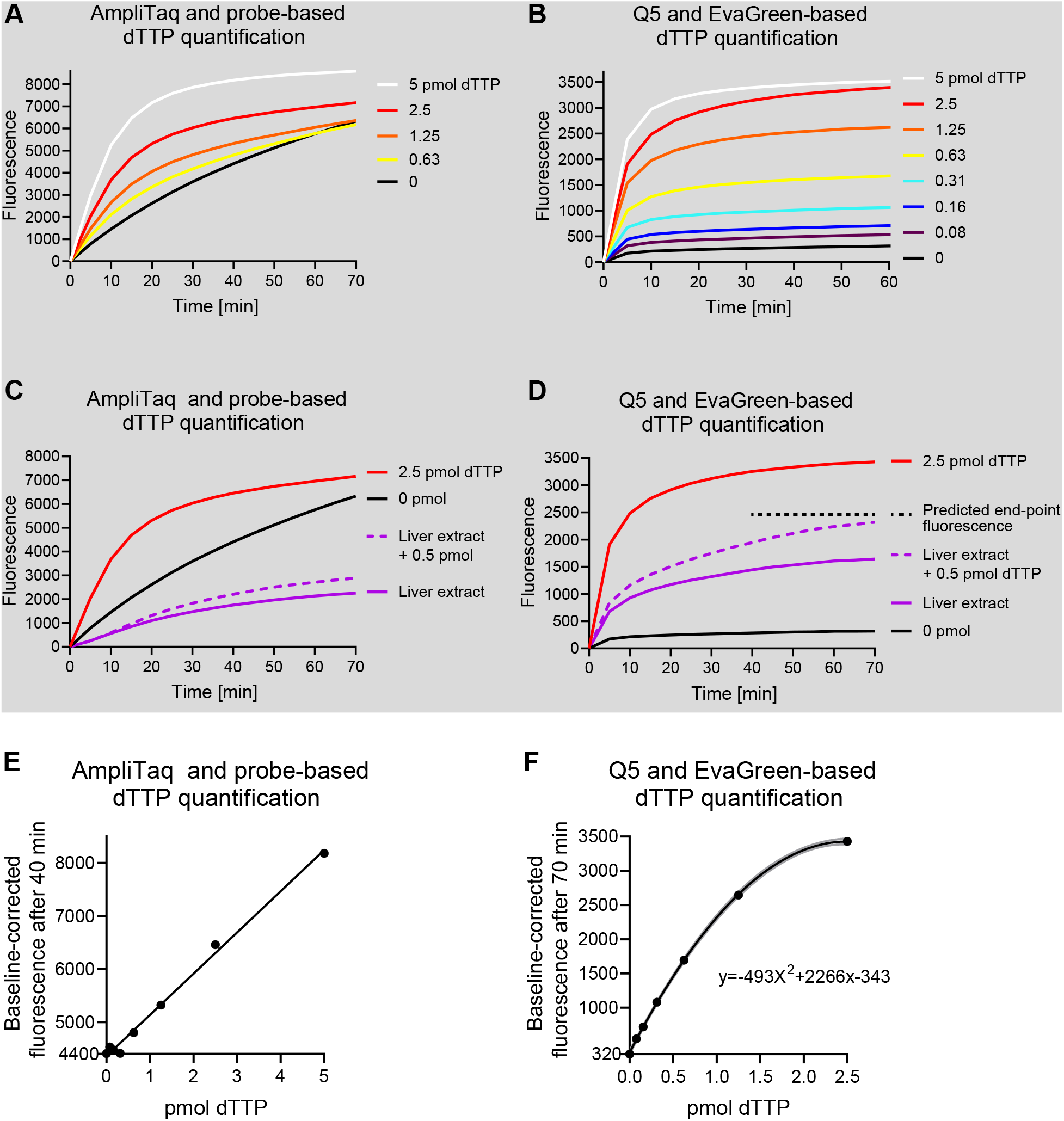
Long synthetic DNA oligonucleotides, an inhibitor-resistant high-fidelity DNA polymerase and EvaGreen detection chemistry allow quantification of dNTPs from mouse liver extracts. (**A**) Representative quantification of dTTP using the published fluorometric probe-hydrolysis-based assay. For clarity, curves from three lowest standard samples (0.31 to 0.08 pmol) are not shown. (**B**) Representative quantification of dTTP using Q5 DNA polymerase, long oligonucleotide template and EvaGreen detection chemistry. (**C**-**D**) dTTP signal generated by the fluorometric methods from mouse liver extracts with and without 0.5 pmol dTTP spike-in calibrant. The final extract volume was diluted to 80 µl per 40 mg of initial tissue weight. (**E**-**F**) Standard curves generated from the end-point baseline-corrected fluorescence values. The lowest y-axis value shows the background signal. Gray lines present the 95% confidence interval of the non-linear curve fit.

**Figure 2.**
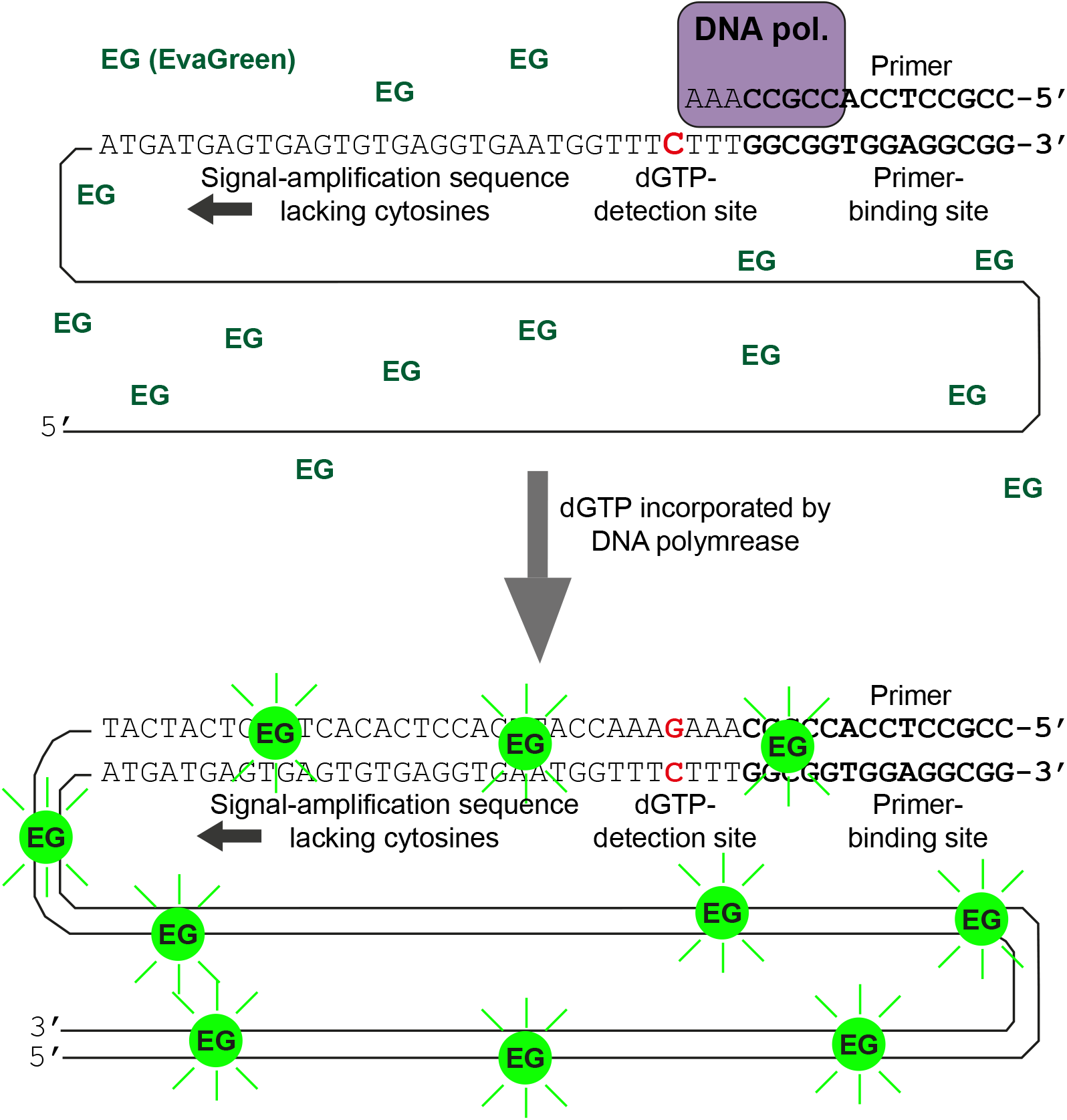
The principle of the novel dNTP assay. Single-stranded DNA templates a comprise 3’ primer-binding site followed by a single detection site for the dNTP to be quantified (here dGTP). The rest of the template, in the 5’ direction, does not contain any cytosines, and its sole purpose is to maximize the amount of double-stranded DNA synthesized per a single dGTP incorporated by the DNA polymerase. The double-stranded DNA binds the EvaGreen fluorochrome and renders it fluorescent, which is measured. In this method, the template length determines the signal amplification.

To test our hypothesis that the error rate of the DNA polymerase could be a major determinant of assay background, we performed the assay with EvaGreen detection using a typical Taq polymerase (AmpliTaq Gold), Phire DNA polymerase (2x fidelity vs Taq), Phusion polymerase (52x fidelity vs Taq) and Q5 polymerase (280x fidelity vs Taq). Indeed, the highest rated fidelity produced the lowest background (Figure 3A). However, using the EvaGreen detection chemistry, the non-proofreading AmpliTaq Gold DNA polymerase also gave a rather stable background in comparison to the probe-based assay, suggesting that unspecific probe hydrolysis could also play a role in the probe-based detection. Our novel EvaGreen-based dNTP assay also worked with Phire and Phusion DNA polymerases without any prior optimization (Figure 3A, B). The Phire DNA polymerase showed somewhat faster reaction kinetics than the other three enzymes. With this DNA polymerase, the reactions were at near completion after 15 minutes whereas 50 – 60 minutes was required for the other enzymes. Based on the lowest background signal, and reported (10) inhibitor resistance in DNA extraction-free PCR, we chose to continue the optimization with Q5 DNA polymerase. This polymerase and its commercial buffer also showed the lowest autofluorescence (Supplementary Figure 3).

**Figure 3.**
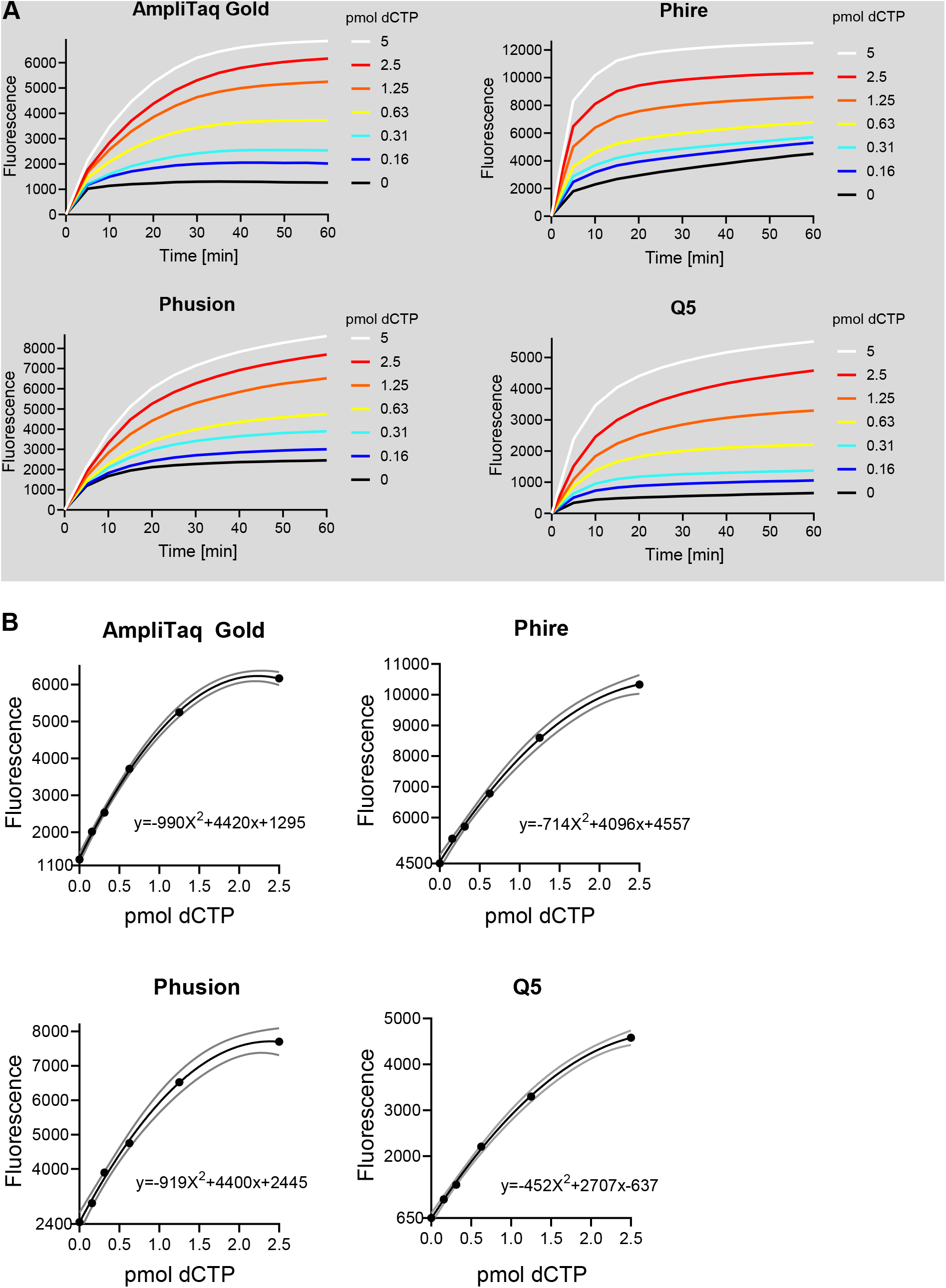
Effect of four different DNA polymerases on the assay kinetics. (**A**) Time-dependent increase in baseline-corrected fluorescence under different concentrations of limiting dNTP (dCTP). (**B**) Standard curves with second-order polynomial curve fit generated from end-point fluorescence values (1h). Gray lines present the 95% confidence interval of the curve fit. Y-axis starts from the background signal. The same primer (0.4 µM), template (0.4 µM), non-limiting dNTP (100 µM) and EvaGreen (1x) concentrations were used for all reactions in buffers supplied by the manufacturers of the DNA polymerases. Reaction temperatures were 60°C for AmpliTaq Gold (17.5 U/ml), 61.7°C for Phire (20 µl/ml) and Phusion (10 U/ml), and 68°C for Q5 (10 U/ml). The reaction temperature for Phire, Phusion and Q5 were chosen based on recommendations by the manufacturer. In the case of AmpliTaq Gold, the reaction temperature was selected based on the published probe-hydrolysis based assay.

### Optimization of the reaction conditions

We systematically optimized the concentration of key reaction components and temperature to maximize sensitivity and to decrease cost of the assay (Figure 4). A decrease in primer and template concentration from 0.4 to 0.2 µM almost halved the background without notable reduction in specific signal (Figure 4A). A slight excess of primer (1.25-fold) over the template marginally improved the assay at reaction temperature of 68°C (Figure 4B). Non-limiting dNTP concentrations between 25 to 100 µM had little effect on the assay performance (Figure 4C). We chose the concentration of 50 µM as optimal for later runs. Polymerase concentration of 20 U/ml showed marginal improvement over 10 U/ml while 5 U/ml compromised the assay (Figure 4D). Therefore, we chose the polymerase concentration of 20 U/ml for maximum sensitivity and 10 U/ml for measurements when the maximum sensitivity is not required. The EvaGreen concentration recommended by the manufacturer for qPCR proved to be optimal (Figure 4E and Supplementary Figure 4). We varied the reaction temperature between 62 and 72°C and found that optima lied between 66 and 68°C (Figure 4F). Lower temperatures increased the background and higher ones compromised the assay.

**Figure 4.**
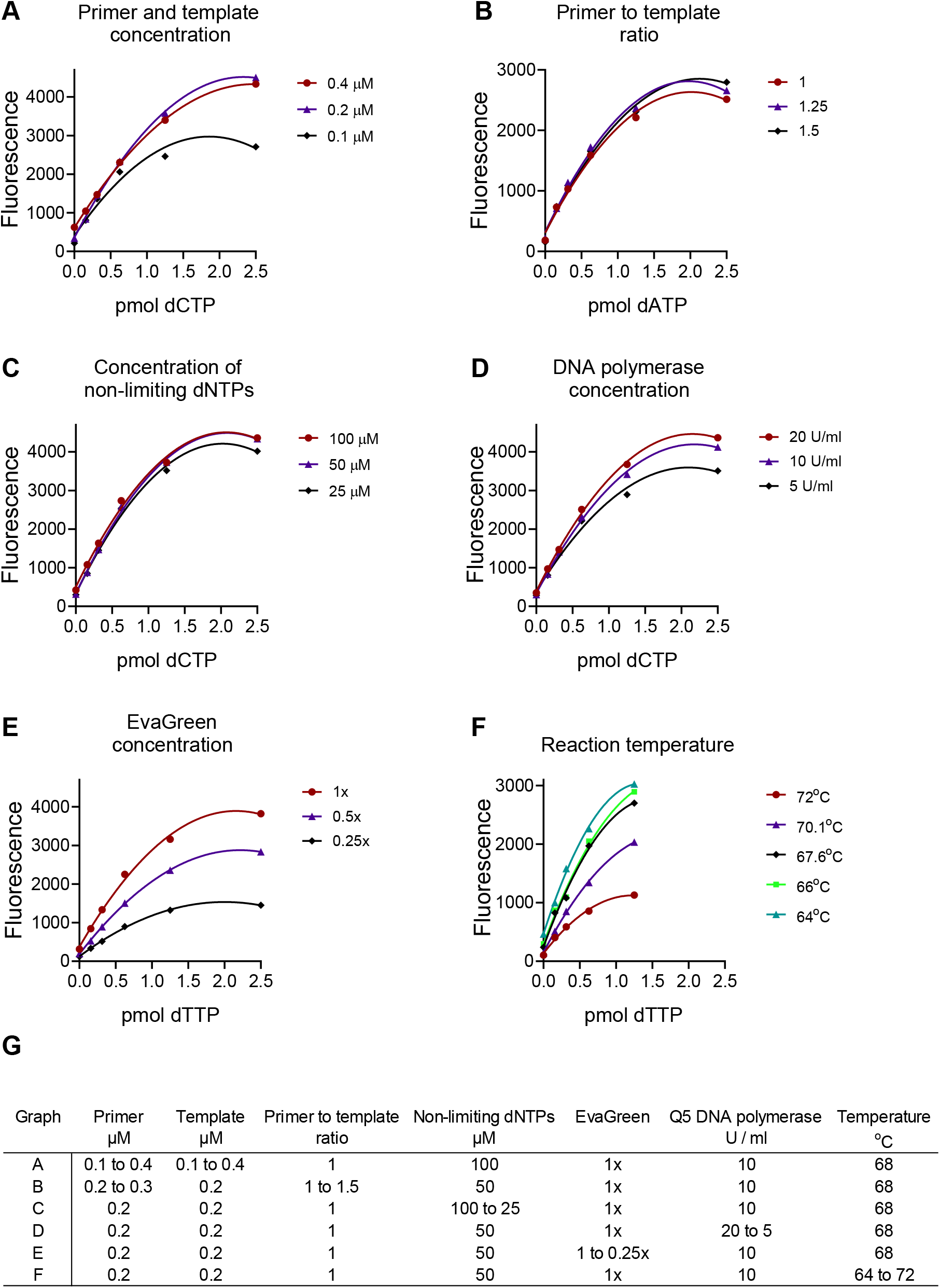
Optimization of EvaGreen- and Q5 DNA polymerase-based dNTP quantification. (**A-F**) Each critical reaction component and reaction temperature were varied and the outcome evaluated. (**G**) Assay conditions in graphs **A** to **F**. The standard curves were generated from end-point (1h) baseline-corrected fluorescence values.

After all optimization, we found that the assay produced a quasilinear response for input of 0.1 to 0.94 pmol (R^2^>0.99) (Figure 5). Second-order polynomial curve fit, however, described the standard curve most accurately and gave a dynamic quantitative range of 0.1 to 2 pmol. The lowest assay concentration we tested was 5.9 nM (0.03 pmol) and signal from this concentration could still be distinguished from the background at signal-to-noise ratios ranging from 3.5 to 5.3.

**Figure 5.**
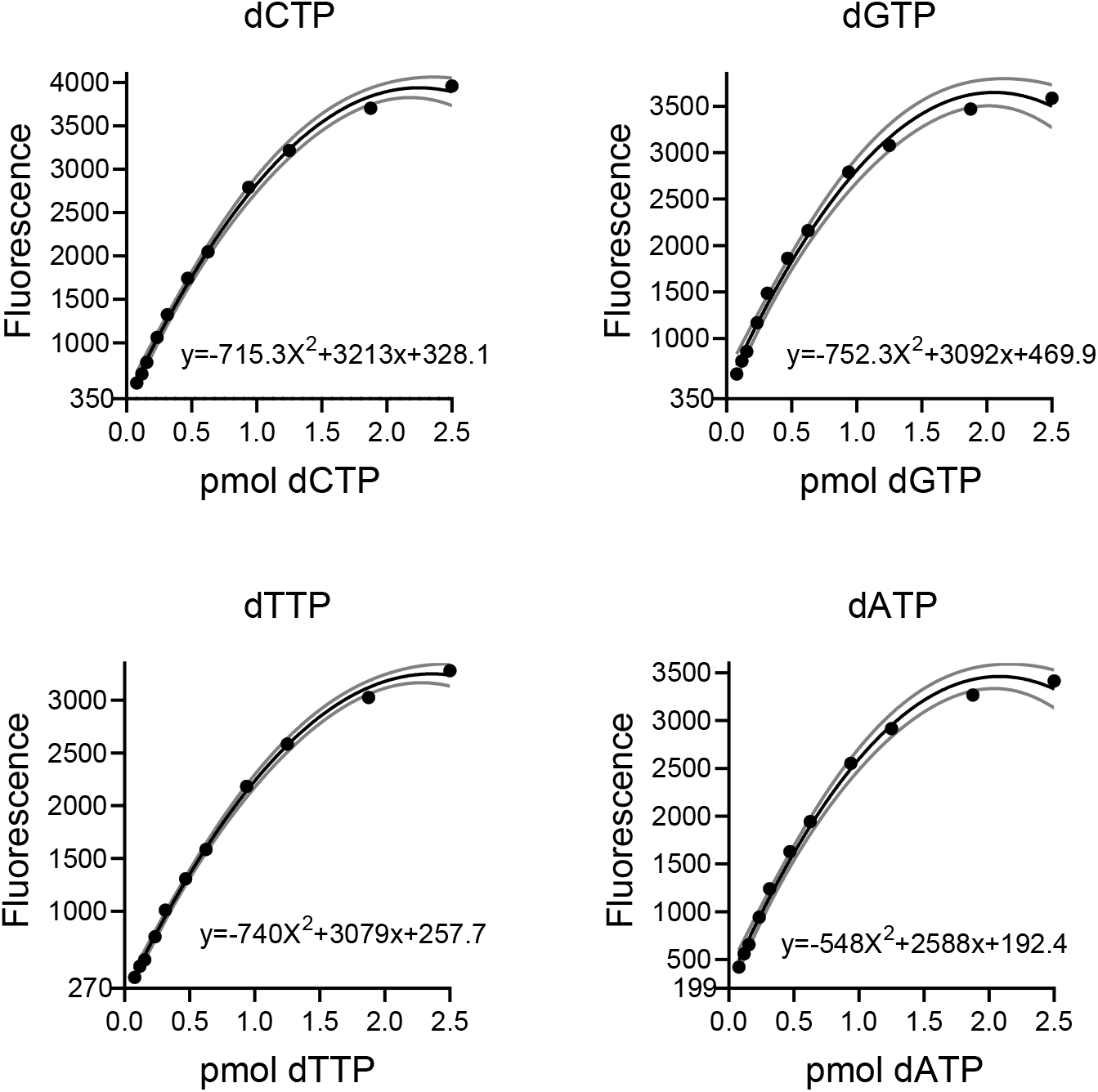
Standard curves after assay optimization. Gray lines present the 95% confidence interval of the second-order polynomial curve fit. The lowest value of y-axis represents the background signal.

### Optimization of sample preparation

As the starting point, we chose the well-validated 60% methanol extraction, which has been shown to give dNTP recoveries higher than 90% (6, 7). We also adopted a 3-kDa cut-off centrifugal filtration from a previous study (7) to remove any remaining DNA and other macromolecules that could increase EvaGreen background fluorescence, and produce unspecific DNA polymerisation, or polymerase inhibition. For enzymatic assays the methanol extracts have typically been evaporated to dryness in vacuum. To minimize nucleotide hydrolysis, oxidation, and deamination reactions, we chose a more rapid approach and extracted the methanol with diethyl ether. The residual diethyl ether was easily removed by boiling the sample tubes cap open for 1 min or by a short evaporation (~15 min) in a vacuum centrifugal evaporator at 65°C. The diethyl ether extraction step was also presumed to remove some putative non-polar polymerase inhibitors such as hemin, bilirubin and bile acids based on the solubility of these compounds.

### Determination of dNTP concentrations in cultured cells

As a biological validation of the novel method, we measured dNTP levels from a mouse hepatoma cell line (Hepa 1-6) cultured with or without 5 µM 5-fluorouracil, a thymidylate synthase inhibitor that blocks dTTP biosynthesis (Table 1). After a 6-hour treatment, 5-fluorouracil had depleted dTTP levels to one fifth of the levels in untreated cells. 5-fluorouracil also decreased dGTP and dCTP concentration to some extent, while that of dATP was increased. We also manipulated dNTP levels by letting the cells grow to a near-confluent monolayer in order to decrease the number of proliferating cells. As expected, the confluent cells had a clear decrease in all four dNTPs (Table 1). Next, we tested the effect of inhibition of pyrimidine biosynthesis by blocking mitochondrial respiratory complex III with 200 nM myxothiazol for 6h. Complex III is required for the activity of dihydroorotate dehydrogenase (DHODH), a mitochondrial inner-membrane quinone oxidoreductase needed for the *de novo* biosynthesis of all pyrimidine nucleotides (11). Instead of resulting in a biased ratio of pyrimidine to purine dNTPs, myxothiazol decreased all dNTPs to levels similar as or lower than in non-cycling (confluent) cells, suggesting that it halted cell proliferation (Table 1). This was dependent on complex III because the myxothiazol-induced decline in dNTP levels was prevented by the expression of Pacific oyster alternative oxidase (PoAOX), an enzyme that can bypass a blockade of the complex III-IV segment of the respiratory electron transfer (Supplementary Figure 1), as also previously shown by its heterologous expression in yeast (8).

**Table 1.** dNTP concentrations in cultured mouse hepatoma cells (Hepa 1-6), pmol/10^6^ cells

| Treatment/condition | Transgene | dTTP | dATP | dCTP | dGTP |
| --- | --- | --- | --- | --- | --- |
| Untreated |  | 42 ± 6.9 | 14.5 ± 2.3 | 133 ± 25 | 4.7 ± 0.8 |
| 5 µM 5-fluorouracil |  | 7.8 ± 0.9 | 24.1 ± 0.9 | 67 ± 7.2 | 1.4 ± 0.1 |
| Confluent cells |  | 4.5 | 2.7 | 5.2 | 2.2 |
| Vehicle (0.008% EtOH) | empty vector | 40.1 ± 2.5 | 17.9 ± 2.4 | 161 ± 13 | 7.5 ± 0.5 |
| 200 nM myxothiazol | empty vector | 3.8 ± 0.6 | 4.6 ± 1.5 | 3.5 ± 0.5 | 0.5 ± 0.4 |
| 200 nM myxothiazol | *poAOX* | 43 ± 4.4 | 22.8 ± 3.8 | 163 ± 30 | 12.6 ± 3.8 |

The values represent mean ± SD. The number of independent replicates is three for all conditions except for confluent cells for which n=1. Treatment time for the experiments was 6h.

### Determination of dNTP concentrations in mouse tissues

To demonstrate the true utility of our method, we set out to measure dNTP pools from mouse liver, heart and skeletal muscle, tissues comprising mostly non-dividing cells and therefore having minimal dNTP concentrations. Our method was easily able to quantify all four dNTPs from these four tissues (Table 2) with detection efficiency values ranging from 90 to 100% (Table 3). Across the four tissues, dATP was the least abundant dNTP (~0.2 pmol/mg tissue) while dTTP and dGTP concentrations were at least 2-fold higher. Heart tissue showed the most asymmetric dNTP pools with dGTP concentration approximately 4 pmol/mg while the concentrations of other dNTPs were more similar to that in liver and skeletal muscle.

**Table 2.**
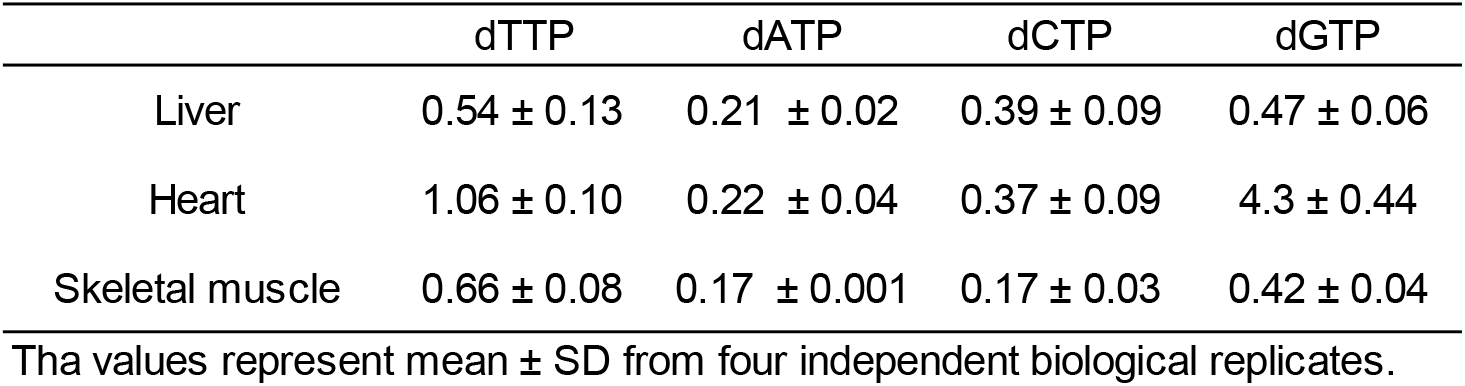
dNTP concentrations in mouse tissues, pmol/mg tissue

**Table 3.** Efficiency of detection reaction (%)

|  | dTTP | dATP | dCTP | dGTP | Extract volume per initial tissue weight |
| --- | --- | --- | --- | --- | --- |
| Liver | 100 | 94 | 95 | 90 | 3 µl / mg |
| Liver | 93 | 96 | 96 | 91 | 4 µl / mg |
| Heart | 96 | 101 | 93 |  | 4 µl / mg |
| Heart |  |  |  | 95 | 28 µl / mg* |
| Skeletal muscle | 99 | 88 | 93 | 89 | 4 µl / mg |

The reaction efficiencies were estimated by performing the measurements with and without a spike-in of purified dNTPs (corresponding to 0.25 pmol / reaction).
\* Heart extracts had to be diluted further for the measurement of dGTP to reach the quantitative range of the assay.

## DISCUSSION

Balanced dNTP pools are vital for efficient error-free DNA synthesis (1). The current knowledge related to regulation and fluctuations in dNTP pools derive mainly from cell culture experiments. Tissue dNTP pools in health and disease have been studied surprisingly little. This has likely been partly because of the low concentrations of dNTPs in tissue samples and the lack of a sensitive-enough methodology. To fill this methodological gap, we developed a simple robust method able to quantify each of the four canonical dNTP from mouse tissue biopsies.

Our novel and the published enzymatic fluorometric assay share a number of benefits over the traditional methods based on radionucleotide incorporation. First of all, the quantification reaction takes place in a single microplate well and requires only pipetting of a reagent mix and the sample. Secondly, a typical qPCR instrument can be used for the temperature control and to read the fluorescence. Whereas in the case of radionucleotide-based detection, the reaction products have to be transferred onto filter paper, washed multiple times, and quantified using a scintillator. Such manual labour-requiring steps are prone to human error and limit the feasible number of samples that can be analysed simultaneously. In addition, handling of radioactive reagents requires permits and stringent safety measures. Recently, an improved enzymatic radionucleotide-based assay for dNTPs was developed in which biotinylated oligonucleotide templates are incorporated onto streptavidin-coated microplates to facilitate washing steps and recovery of the synthetized radio-labelled DNA (12). The inventors of this assay measured dNTPs from several cell lines, but they did not report suitability of the assay for tissue samples. Nevertheless, our novel assay outperforms published enzymatic assays in simplicity, cost effectiveness, and the sensitivity is similar or better.

A novel innovation of our method is the utilization of the EvaGreen dye and recent advances in chemical oligonucleotide synthesis that allow production of ultralong (~200 nt) ssDNA templates at reasonable cost. EvaGreen is a well-characterized intercalating DNA fluorochrome that shows higher specificity for double-stranded versus single-stranded DNA than the more traditional SYBR Green fluorochrome used in qPCR (13). Moreover, it can be used at saturating concentration without DNA polymerase inhibition. The fluorescence of EvaGreen is proportional to dsDNA concentration and length, and therefore the template length sets the signal amplification in our method. One benefit of EvaGreen-based detection as compared to fluorescent probes is that the choice of DNA polymerase is not limited to enzymes possessing 5’ to 3’ exonuclease activity to hydrolyse the probe. An appropriate DNA polymerase is crucial for accurate quantification of dNTPs by the enzymatic method. For instance, the commonly used Klenow fragment has been shown to substitute corresponding ribonucleotide for the dNTP to be quantified (5). Taq polymerases can better discriminate deoxyribonucleotides from ribonucleotides but, similar to the Klenow fragment, are able to incorporate dUTP instead of dTTP (7). In addition, in the case of tissue extracts, the DNA polymerase has to be able to tolerate a considerable amount of inhibitory impurities. To our knowledge, the utility of modern inhibitor-resistant high-fidelity DNA polymerases in enzymatic determination of dNTPs has not been investigated before, even though highly unbalanced dNTPs required by the assays are expected to lead to eventual incorporation of wrong deoxyribonucleotides with concomitant elevation of background signal. We, indeed, found that the DNA polymerase with the highest rated fidelity gave the lowest background, but polymerase error-rate was clearly not the sole factor determining the background. In fact, we found that EvaGreen-based quantification robustly worked with four different DNA polymerases without any prior optimization. Importantly, after optimization, we could reach reliable quantification down to 100 fmol (20 nM) with a detection limit being lower than 30 fmol (3 nM). As a comparison, a radionucleotide-based method has been reported to reach a 100 fmol quantification limit (data not shown and criteria for LLOQ not reported) (5). The most sensitive published HPLC-MS system has a LLOQ of 60 fmol, and typically the LLOQ values for chromatographic methods range from 0.25 to 10 pmol (6). The previous fluorometric assay based on double-quenched probes had LLOQ values ranging from 0.8 to 1.3 pmol (7). Less stringent criteria (5 SD over background) for LLOQ was, however, used by these authors than by us. Recently, a rather elaborate enzymatic two-stage dNTP detection involving a fluorescent probe, two DNA fragments, two different DNA polymerases and a DNA ligase was developed (14). This assay could reach a detection limit that is orders of magnitude lower than any other existing method could. However, this complex assay was developed for single-molecule DNA sequencing and its utility has not been demonstrated with biological samples.

The main reason we set out to develop a new method was the need to quantify minute dNTP concentrations from reasonably small mouse tissue biopsies. The first enzymatic assay for dNTPs was published 1969 (15). Since then, small improvements to increase sensitivity and specificity have been reported (4, 5, 12, 16). In all these studies, however, only extracts from cultured cells were used as a biological sample. Yet, enzymatic methods have been used to measure dNTP concentrations from liver (17) and skeletal muscle samples (18–20), but no method validation data for tissue samples has been reported in these studies. In skeletal muscle samples, individual dNTP concentrations have ranged from less than 0.02 to 0.3 pmol/mg with the biggest variation seen in dATP levels (18, 19). Interestingly, a HPLC-MS-based method has also shown surprisingly low dATP concentrations for mouse heart tissue (0.02 pmol/mg) (6). Although the ribonucleotide ATP is the physiological substrate for cellular ATPases, at least some of these enzymes can use dATP instead of ATP (21). We speculate that the variation in dATP levels could be related to ischemia during sample collection and delayed quenching of enzymatic activity. Similar to two chromatographic quantifications (6, 22), we found heart dGTP concentration to be 5 to 15-fold higher than that of other three dNTPs. Mitochondrial dNTP pools are extremely asymmetric with dGTP concentration being more than 10-fold higher than the concentration of other dNTPs (23). This asymmetry was reported to be especially notable in heart mitochondria (155-fold excess dGTP) (23). Therefore, it is obvious that in mitochondria-rich tissues such as in heart, dGTP would be the most abundant dNTP. In summary, according to our results and the published literature the tissue dNTP concentrations are generally low; only a fraction of a picomole per mg of tissue.

We optimized key parameters of the dNTP assay for Q5 DNA polymerase. Still, more extensive optimization could result in even higher sensitivity. Higher signal amplification could, for example, be achieved by increasing the template length even further. Dynamic range could be adjusted by varying the number of dNTP detection sites in the template, as done for the probe-based assay (7), or using a higher template concentration. Our assay could also be modified to quantify dUTP by comparing results from runs performed using a DNA polymerase that can and another that cannot incorporate dUTP. We acknowledge the following limitations of our assay. First, it is necessary to test new sample (e.g. animal tissue) types for the presence of interfering compounds by performing a dilution series or spiking the samples with known amounts of dNTPs. When we diluted the extracts as instructed here, and the reactions were allowed to proceed to near completion, the sample-related interference was insignificant. However, we found that too concentrated extracts can lead to underestimation of dNTP concentrations. The assay produces a standard curve that follows second-order polynomial equation. Thus, if inhibition is present, the classical Standard Addition Method cannot be used as a correction. Tissue extracts tend to show some autofluorescence. Therefore, we found it absolutely necessary to subtract baseline fluorescence from the end-point fluorescence values. This correction also decreased well-to-well variation. An approximate cost of our assay, excluding sample preparation, is 34 EUR (37 USD) per 100 detection reactions.

In summary, we developed an enzymatic fluorometric assay that is simple, inexpensive, and most importantly has sensitivity and robustness to measure dNTPs from non-proliferating cells or from tissue extracts. When technical triplicates are used for the measurements, as little as 15 mg tissue or 0.5×10^6^ cultured cells is sufficient. Moreover, 96 and 384-well formats of the assay allow a large number of samples to be measured simultaneously. We have analysed more than 40 tissue samples in triplicates using our method in 384-well format (results to be published elsewhere). Only handful of studies have addressed the topic of tissue dNTP levels in health and disease (6, 17–19). As nucleotide metabolism is more and more studied in the field of cancer biology and in relation to metabolic diseases such as mitochondrial disorders (19, 24) the availability of an easily applicable dNTP assay for use in the general laboratory is of the utmost importance.

## Supporting information

Step-by-step protocol

## ACKNOWLEDGEMENTS

We thank Elisa Alppila for technical assistance.

## FUNDING

This work was supported by grants from Finnish Physicians’ Society (to V.F.), Foundation for Pediatric Research in Finland (to V.F.), Medicinska Understödsföreningen Liv och Hälsa (to V.F.), Folkhälsan Research Center (to V.F., J.K.), Magnus Ehrnrooth Foundation (to V.F. and R.B.) and Biomedicum Helsinki Foundation (to J.P.).

