## Supplementary Figures and Tables for "A sensitive assay for dNTPs based on long synthetic oligonucleotides, EvaGreen dye, and inhibitor-resistant high-fidelity DNA polymerase"

**Supplementary Table 1**. DNA oligonucleotides used in dNTP assays

| | Sequences in 5' to 3' direction |
|---|---|
| **EvaGreen-based detection** | |
| Detection primer | CCGCCTCCACCGCC |
| dTTP detection template | TCGCTCGCTCTTGCCTCGGTCCTCGCTCGCTCTTGCCTCGGTCCTCGCTCGCTCTTGCCTCGGTCCTCGCTCGCTCTTGCCTCGGTCCTCGCTCGCTCTTGCCTCGGTCCTCGCTCGCTCTTGCCTCGGTCCTCGCTCGCTCTTGCCTCGGTCCTCGCTCGCTCTTGCCTCGGTCCTTTATTT**GGCGGTGGAGGCGG** |
| dATP detection template | AGACAGACACAAGACACAGACCAGACAGACACAAGACACAGACCAGACAGACACAAGACACAGACCAGACAGACACAAGACACAGACCAGACAGACACAAGACACAGACCAGACAGACACAAGACACAGACCAGACAGACACAAGACACAGACCAGAGAGACACAACAGACGGAGGAAATAAA**GGCGGTGGAGGCGG** |
| dCTP detection template | CCACTCACTCTTACCTCAATCCCCACTCACTCTTACCTCAATCCCCACTCACTCTTACCTCAATCCCCACTCACTCTTACCTCAATCCCCACTCACTCTTACCTCAATCCCCACTCACTCTTACCTCAATCCCCACTCACTCTTACCTCAATCCCCACTCACTCTTACCTCAATCCTTTGTTT**GGCGGTGGAGGCGG** |
| dGTP detection template | GGAGTGAGTGTGAGGTGAATGATGAGTGAGTGTGAGGTGAATGTAGAGTGAGTGTGAGGTGAATGATGAGTGAGTGTGAGGTGAATGTAGAGTGAGTGTGAGGTGAATGATGAGTGAGTGTGAGGTGAATGTAGAGTGAGTGTGAGGTGAATGATGAGTGAGTGTGAGGTGAATGGTTTCTTT**GGCGGTGGAGGCGG** |
| **Probe-based detection** | |
| Detection primer | CCGCCTCCACCGCC |
| dTTP probe | **FAM**/AGGACCGAG/**ZEN**/GCAAGAGCGAGCGA/**IBFQ** |
| dATP probe | **FAM**/TGGTCCGTG/**ZEN**/GCTTGTGCGTGCGT/**IBFQ** |
| dCTP probe | **FAM**/AGGATTGAG/**ZEN**/GTAAGAGTGAGTGG/**IBFQ** |
| dGTP probe | **FAM**/ACCATTCAC/**ZEN**/CTCACACTCACTCC/**IBFQ** |
| dTTP detection template | **TCGCTCGCTCTTGCCTCGGTCC**TTTATTT**GGCGGTGGAGGCGG** |
| dATP detection template | **ACGCACGCACAAGCCACGGACC**AAATAAA**GGCGGTGGAGGCGG** |
| dCTP detection template | **CCACTCACTCTTACCTCAATCC**TTTGTTT**GGCGGTGGAGGCGG** |
| dGTP detection template | **GGAGTGAGTGTGAGGTGAATGG**TTTCTTT**GGCGGTGGAGGCGG** |

The pimer-binding sites are shown as bolded black letters at 3' end. Red letters shows the locations of dNTP-detection sites. The probe-binding sites are marked with blue letters.
FAM, 5(6)-carboxyfluorescein
ZEN, a proprietary quencher by Integrated DNA Technologies
IBFQ (Iowa Black FQ), a proprietary quencher by Integrated DNA Technologies

Supplementary Table 2. Critical reagents, materials and intrumentation

| | Manufacturer | Catalogue number |
|---|---|---|
| **Reagents** | | |
| MeOH (analytical grade) | Fisher Scientific | M/4000/PC17X |
| Stabilized diethyl ether (less than 1 year old) | Sigma | 296082 |
| 100 mM dNTP solutions | Thermo Fisher | R0181 |
| 20x EvaGreen | Biotium | 31000 |
| Q5 High-Fidelity DNA polymerase* | New England Biolabs | M0491 |
| Q5 Hot Start High-Fidelity DNA polymerase* | New England Biolabs | M0493 |
| AmpliTaq Gold DNA polymerase | Thermo Fisher | N8080241 |
| Phire Hot Start DNA II DNA polymerase | Thermo Fisher | F122 |
| Phusion High-Fidelity DNA polymerase | Thermo Fisher | F530 |
| **Materials** | | |
| Amicon Ultra-0.5 Centrifugal Filter Unit, 3 kDa | Merck | UFC500396 |
| Hard-Shell® 384-Well PCR Plates, thin wall, skirted, black/white | Bio-Rad | HSP3865 |
| Microseal 'B' PCR Plate Sealing Film | Bio-Rad | MSB1001 |
| **Instrumentation** | | |
| Speed-Vac Plus SC110A evaporator | Savant Instruments | |
| CFX384 qPCR instrument | Bio-Rad | |

DNA oligonucleotides are listed in Supplementary Table 1.

* No difference in performance was observed between these two DNA polymerase preparations.

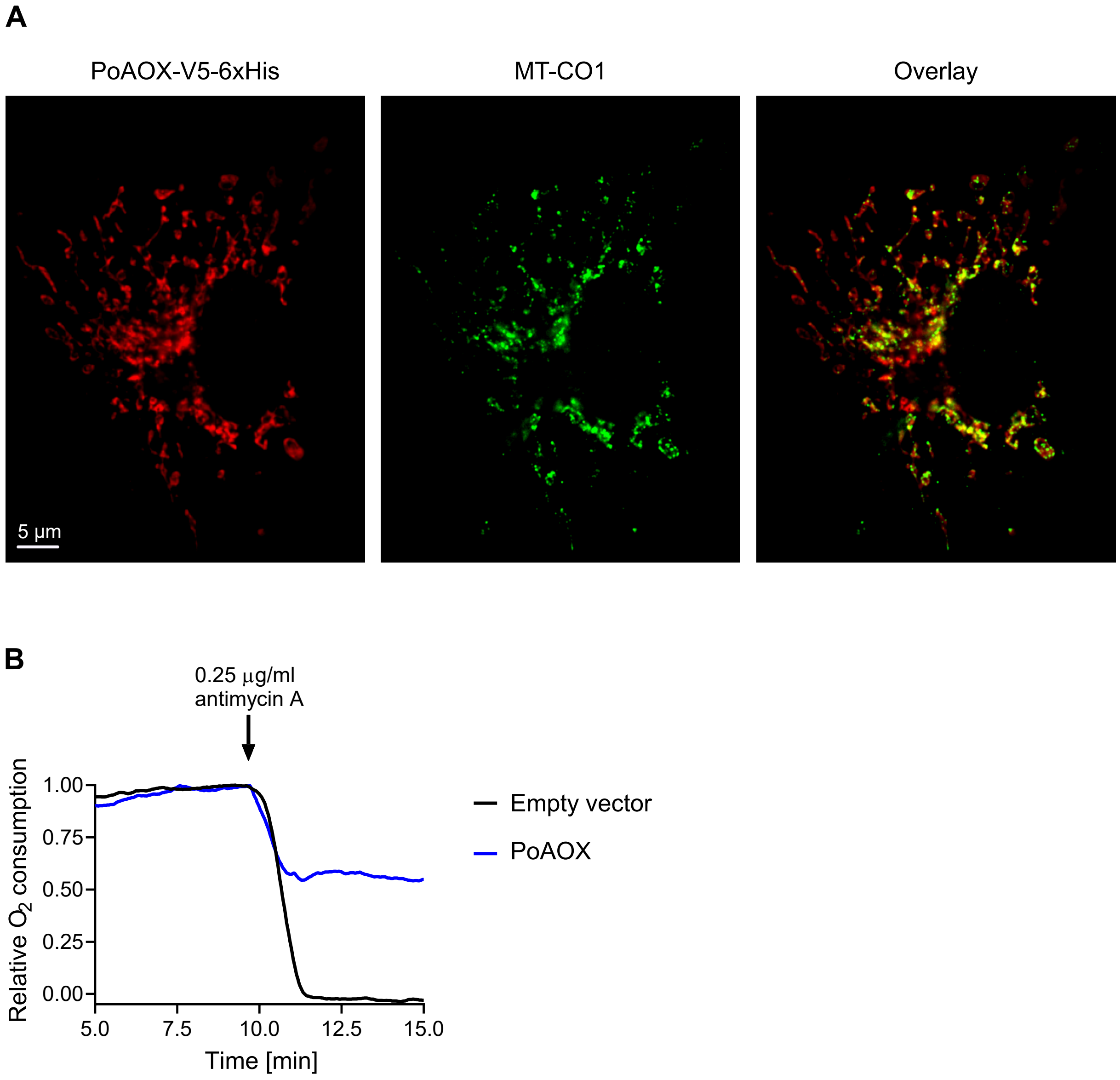


**Supplementary Figure 1.** Pacific oyster (*Crassostrea gigas*) alternative oxidase (poAOX) localizes to mitochondria and confers antimycin A-resistant respiration in mammalian cells. (**A**) Transfected COS-1 cells immunostained for V5-tagged PoAOX and the endogenous mitochondrial protein MT-CO1 (cytochrome c oxidase subunit 1). (**B**) Respiration by intact Hepa1-6 cells in complete DMEM at +37°C was measured using Oxygraph-2k (OROBOROS instruments). Alternative oxidase activation was assessed by the addition of the respiratory complex III inhibitor antimycin A. Residual oxygen consumption after the subsequent addition of rotenone was set as background. Maximal basal respiration was set as another reference state and the data were scaled according to these two reference states.

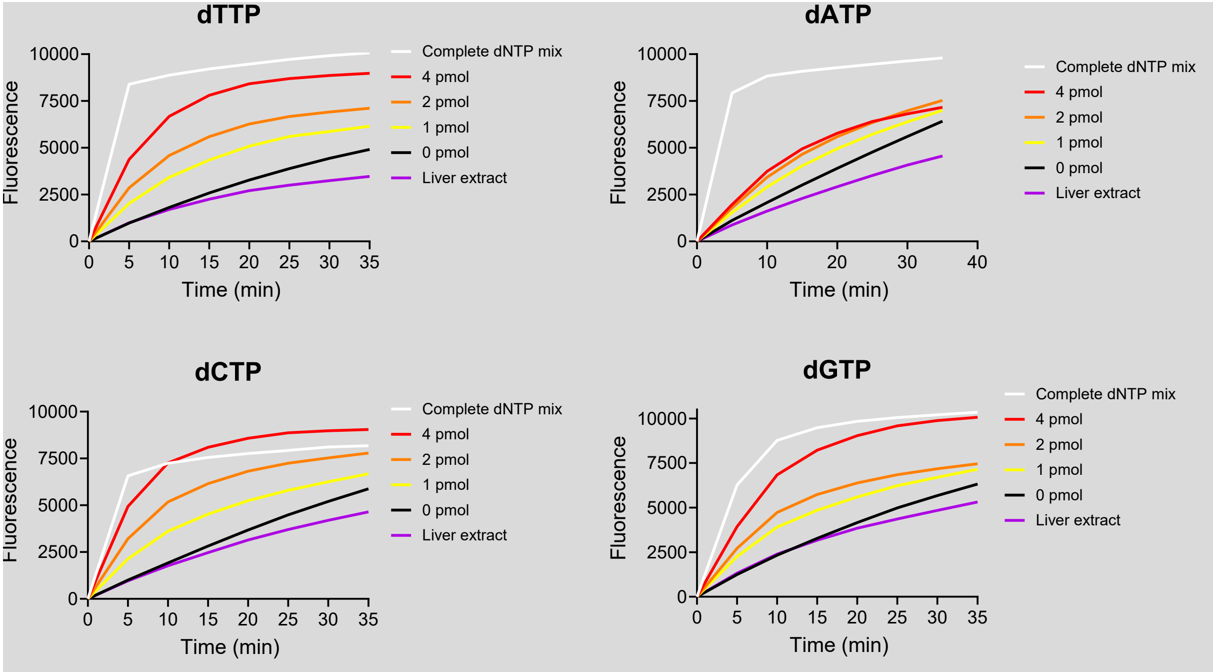


**Supplementary Figure 2.** Suitability of AmpliTaq Gold and probe hydrolysis-based dNTP quantification for liver extracts. Here, we followed a published protocol to assess dNTPs from liver extracts. Even dilute liver extracts (12 µl/mg of initial tissue weight) interfered with the assay leading to signal that was lower than the background.

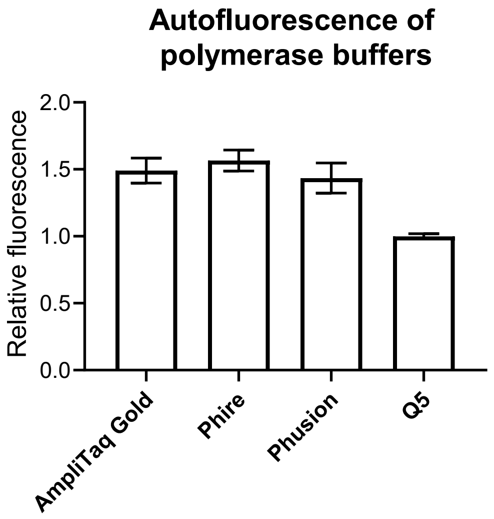


**Supplementary Figure 3.** Autofluorescence of tested DNA polymerase buffers.

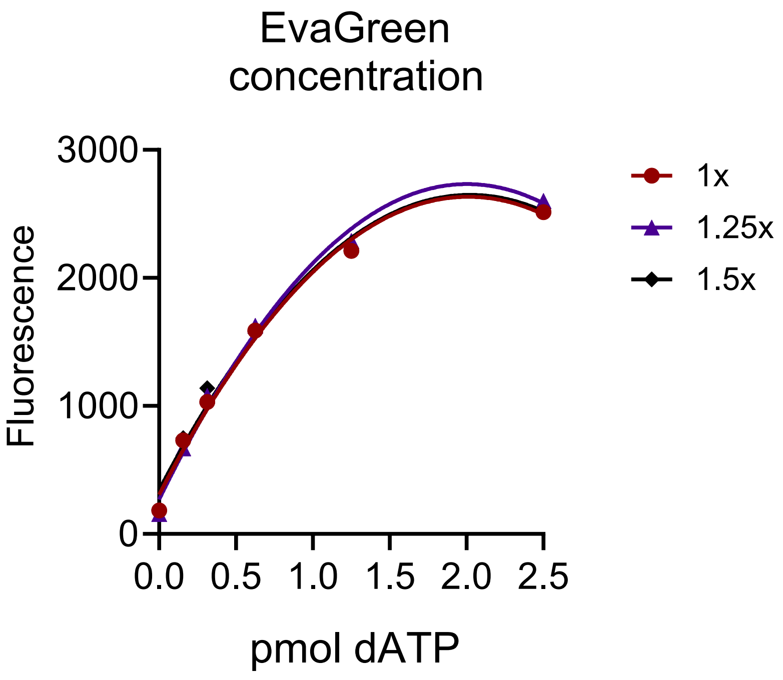


**Supplementary Figure 4.** Increasing the EvaGreen concentration above the manufacturer's recommended concentration does not benefit the assay.
