## Supplementary material for "A sensitive assay for dNTPs based on long synthetic oligonucleotides, EvaGreen dye, and inhibitor-resistant high-fidelity DNA polymerase": Step-by-step protocol

### Step-by-step protocol for measurement of dNTP concentrations in mouse tissues

#### Reagents and materials

- Ice-cold 60% MeOH

- Amicon Ultra-0.5-ml centrifugal filters, 3kDa (Merck, UFC500396). Mark and weight the collection tubes before use. Equilibrate with 60% MeOH before use. Do not let dry.

- Diethyl ether (CAUTION! Intoxicating, flammable and explosive. Open diethyl ether-containing bottles and vials only in a fume hood. Check the expiry date. Do not use if expired.)

- DNA oligonucleotides

Sequences in 5' to 3' direction:

Nucleotide detection primer: CCGCCTCCACCGCC

dTTP_template_197nt
TCGCTCGCTCTTGCCTCGGTCCTCGCTCGCTCTTGCCTCGGTCCTCGCTCGCTCTTGCCTCGGTCCTCGCTCGCTCTTGC
CTCGGTCCTCGCTCGCTCTTGCCTCGGTCCTCGCTCGCTCTTGCCTCGGTCCTCGCTCGCTCTTGCCTCGGTCCTCGCTC
GCTCTTGCCTCGGTCCTTT**A**TTT**GGCGGTGGAGGCGG**

dATP_template_197nt

AGACAGACACAAGACACAGACCAGACAGACACAAGACACAGACCAGACAGACACAAGACACAGACCAGACAGACACAAGA
CACAGACCAGACAGACACAAGACACAGACCAGACAGACACAAGACACAGACCAGACAGACACAAGACACAGACCAGAGAG
ACACAACAGACGGAGGAAA**T**AAA**GGCGGTGGAGGCGG**

dCTP_template_197nt

CCACTCACTCTTACCTCAATCCCCACTCACTCTTACCTCAATCCCCACTCACTCTTACCTCAATCCCCACTCACTCTTAC
CTCAATCCCCACTCACTCTTACCTCAATCCCCACTCACTCTTACCTCAATCCCCACTCACTCTTACCTCAATCCCCACTC
ACTCTTACCTCAATCCTTT**G**TTT**GGCGGTGGAGGCGG**

dGTP_template_197nt

GGAGTGAGTGTGAGGTGAATGATGAGTGAGTGTGAGGTGAATGTAGAGTGAGTGTGAGGTGAATGATGAGTGAGTGTGAG
GTGAATGTAGAGTGAGTGTGAGGTGAATGATGAGTGAGTGTGAGGTGAATGTAGAGTGAGTGTGAGGTGAATGATGAGTG
AGTGTGAGGTGAATGGTTT**C**TTT**GGCGGTGGAGGCGG**

Dissolve the templates to a concentration of 10 µM (e.g. 2.5 nmol DNA and 250 µl $H_2O$). Dissolve the primer to a concentration of 100 µM, and from this prepare a working solution of 10 µM. Store the DNA oligonucleotides at -20ºC.

- NEB Q5 Hot-Start High-Fidelity DNA polymerase

- dNTP set 100 mM Solutions (e.g. Thermo R0181)

- 2.5 mM dNTP mixes:
  dGTP + dCTP + dTTP mix for dATP detection:
  5 µl of 100 mM dGTP
  5 µl of 100 mM dCTP
  5 µl of 100 mM dTTP
  185 µl of $H_2O$

  dCTP + dTTP+ dATP mix for dGTP detection:
  5 µl of 100 mM dCTP
  5 µl of 100 mM dTTP
  5 µl of 100 mM dATP
  185 µl of $H_2O$

  dGTP + dTTP+ dATP+ mix for dCTP detection:
  5 µl of 100 mM dGTP
  5 µl of 100 mM dTTP
  5 µl of 100 mM dATP
  185 µl of $H_2O$

  dGTP + dCTP+ dATP+ mix for dTTP detection:
  5 µl of 100 mM dGTP
  5 µl of 100 mM dCTP
  5 µl of 100 mM dATP
  185 µl of $H_2O$

- dNTP standards: Seven-point 1:2 serial dilution covering 500 to 7.81 nM. This can be prepared from individual dNTP stocks or from a standard 10 mM dNTP mix.

**dNTP extraction from mouse tissues**

Tissue samples should be snap-frozen as fast as possible after cervical dislocation or terminal anaesthesia of the mice. The recommended amount of tissue is 20-40 mg for liver.

1. Homogenize tissue in 550 µl of ice-cold 60% MeOH (up to 40 mg of tissue).

   Liver and other soft tissues:
   First completely crush the tissue using a microtube pestle. Then use a battery-operated microtube pestle homogenizer for 30s.

   Fibrous tissues such as heart and skeletal muscle:
   Mince the tissue into small pieces in ice-cold 60% MeOH using sharp scissors. Transfer the tissue pieces into a roughened glass-to-glass potter tissue grinder. Homogenize using 15 rotating strokes or until the solution is fully homogenous [Note 1].

2. Incubate 3 min. at 95ºC. Cool down on ice.
3. Centrifuge 18500g for 6 min. at +4ºC.
4. Collect supernatant (max 550 µl) into an equilibrated Amicon Ultra-0.5-ml centrifugal filter. The collection tube has to be pre-weighted.
5. Centrifuge at full speed for 60 min. at +4ºC and save the flow-through in the collection tube. (A volume of ~50 µl will remain in the column below the filter line.)
6. Add 1.4 ml ice-cold diethyl ether to the collection tube. Vortex 10s and then shake vigorously for 30s. [Note 2]
7. Spin quickly at 14000g. Remove most of the upper layer with a pipette and discard.
8. Repeat steps 6 and 7.
9. Concentrate the sample and evaporate residual diethyl ether using Speed-Vac: High setting (65ºC) for 15 min.
10. Determine the amount of remaining solution by weighing the tubes (1 mg = 1 µl). Adjust sample volume to 160 µl per 40 mg initial tissue weight by adding $H_2O$.
11. Store the extracts at -80ºC.

**dNTP measurement procedure**

Prepare a master mix for the dNTP to be quantified (dATP, dGTP, dCTP or dTTP). Equilibrate the master mix containing all components except the polymerase to room temperature. Add the polymerase to the master mix immediately before use.

<u>2x master mix recipe for 100 reactions with final volume of 10 µl:</u>

175 µl of $H_2O$
200 µl of 5X Q5 reaction buffer
25 µl of 10 µM nucleotide detection primer
20 µl of 10 µM template
20 µl of 2.5 mM dNTP mix without the dNTP to be quantified
50 µl of 20X EvaGreen stock
10 µl of 2000 U/ ml Q5® High-Fidelity DNA Polymerase

<u>Final concentrations</u>:
Primer, 0.25 µM
Template, 0.2 µM
Non-limiting dNTPs, 50 µM
1 X EvaGreen
Q5 polymerase, 20 U/ml

Assay the samples in triplicates in a 384-well qPCR plate with white wells.

1. Place the qPCR plate on ice.
2. Pipet in reverse mode 5 µl of master mix into the wells.
3. Pipet 5 µl of sample or standards.
4. Program a qPCR instrument to perform the following steps:
   Step 1: 10s at 98ºC.
   Step 2: 1s at 67ºC
   Step 3: read fluorescence.
   Step 4: 5 min at 67ºC

   Repeat step 2 to 4 for 15 cycles (1 h 15 min). [Notes 3 and 4]

5. Turn off the automatic baseline correction by the instrument. Export the raw fluorescence values for data analysis.
6. Generate a standard curve using a second-order polynomial curve fit from the baseline-corrected fluorescence values of the standard samples. Interpolate the concentrations of the samples.

[Note 1] Incomplete homogenization will lead to a severe underestimation of dNTP pools in muscle tissues.

[Note 2] Diethyl ether extracts methanol and hydrophobic metabolites (free heme, bilirubin etc.)

[Note 3] Primer annealing temperature is several degrees higher in Q5 reaction buffer than in typical PCR buffers.

[Note 4] The first fluorescence measurement gives the baseline, which should be subtracted from end-point fluorescence. The reaction should be allowed to proceed for at least 1 h.
